# Geographic patterns of koala retrovirus genetic diversity, endogenization and subtype distributions

**DOI:** 10.1101/2021.11.17.469066

**Authors:** Michaela D. J. Blyton, Paul Young, Ben D. Moore, Keith Chappell

## Abstract

Koala retrovirus subtype A is the youngest endogenized retrovirus, providing a unique system to elucidate retroviral-host co-evolution. We characterised KoRV geography using faecal DNA from 192 samples across 20 populations throughout the koala’s range. We reveal an abrupt change in KoRV genetics and incidence at the Victoria/NSW state border. In northern koalas, *pol* gene copies were ubiquitously present at greater than 5 per-cell, consistent with endogenous KoRV. In southern koalas, *pol* copies were detected in only 25.8% of koalas and always at copy numbers less than one, while the *env* gene was detected in all animals and in a majority at copy numbers of greater than one per-cell. These results suggest that southern koalas carry partial endogenous KoRV-like sequences. Deep sequencing of the *env* hypervariable region revealed three putatively endogenous KoRV-A sequences in northern koalas and a single, distinct sequence present in all southern koalas. Among northern populations, *env* sequence diversity decreased with distance from the equator, suggesting infectious KoRV-A invaded the koala genome in northern Australia and then spread south. The previously described exogenous KoRV subtypes (B-K), two novel subtypes (L and M), and intermediate or hybrid subtypes were detected in all northern koala populations but strikingly absent from all southern animals tested. Apart from KoRV-D, these exogenous subtypes were generally locally prevalent but geographically restricted, producing KoRV genetic differentiation among northern populations. This suggests that sporadic evolution and local transmission of the exogenous subtypes has occurred within northern Australia, but this has not extended into animals within southern Australia.

**Author Summary:** Retrovirus infection is generally synonymous with disease; however, retroviruses can also become endogenous (incorporated into the germline) and thereby directly contribute to the genetic makeup of a species. This has occurred in all vertebrates, yet little is known about the endogenization process. As the youngest virus known to be endogenized, koala retrovirus (KoRV) offers a unique opportunity to study these early stages of co-evolution. This study reveals a comprehensive picture of KoRV biogeography that informs our understanding of how host population history, host suppression and transmission dynamics can influence retroviral evolution. KoRV is also associated with chlamydiosis and neoplasia in the vulnerable koala. Our improved understanding of how KoRV variants are distributed should guide conservation management to help limit disease.

## Introduction

Retroviruses play a unique role in host evolution and health. Generally, retroviruses are transmitted between individuals within a species and cause severe disease through exogenous infection of somatic cells, such as those of the immune system. Some retroviruses, however, can also infect the germline and become a fundamental component of the host’s genome that is thereafter passed onto progeny, a process known as endogenization. Consequently, retroviral genetic sequences have accumulated in the genomes of all vertebrates throughout their evolution and in humans, now comprise 8% of the genome [1]. However, little is known about the early processes by which retroviral sequences colonise a host genome and how this impacts the host and virus. This is because the initial endogenization of most retroviruses occurred many millions of years ago, with the koala retrovirus (KoRV) being an exception [2].

KoRV is a gammaretrovirus that has been associated with chlamydiosis and may cause neoplasia in its solitary arboreal host, the koala (*Phascolarctos cinerus*, Marsupialia) [3–6]. It is thought that KoRV subtype A entered koala populations from an unknown source [2, 7], with endogenization subsequently occurring between 50,000, and 120 years ago [2, 8]. This initial endogenization event is the youngest known and KoRV is one of only a few retroviruses known to exist in both endogenous and exogenous forms [7, 9–12]. As such, KoRV provides a valuable system to elucidate retroviral-host co-evolution. In this study we investigate the biogeography of KoRV across the koala’s natural range to reveal patterns of genetic diversification and differentiation that shed light on exogenous and endogenous adaptation and transmission dynamics.

KoRV infects koalas throughout their natural range in eastern Australia, yet the endogenous and exogenous forms show distinct distributions. In the northern part of the koala’s range (in the states of Queensland (QLD) and New South Wales (NSW)) endogenous KoRV (enKoRV) loci are present in 100% of northern koalas, while in southern Australia (States of Victoria (Vic) and South Australia (SA)) quantitative PCR of the *pol* proviral gene suggests that KoRV only occurs as an exogenous virus at a prevalence of approximately 15 – 50% [6, 13, 14]. It has been suggested that this geographic pattern reflects the spread of endogenous KoRV from the north, where an original endogenization event occurred, to the south [13]. Such a spread would be expected to produce an endogenization gradient. However, the incidence of koalas carrying endogenous KoRV appears to change abruptly between Victoria and NSW [13–15]. No populations containing both koalas with and without endogenous KoRV have been reported. An alternative explanation proposed is that koalas in southern Australia are resistant to KoRV and/or have eliminated full length, replication competent KoRV from their genomes [16]. Some koalas in SA express the terminal regions of the KoRV genome [16], which could indicate they carry either degraded KoRV variants and/or a apparently replication-incompetent, recombination derived KoRV-like element known as recKoRV [17]. A key factor is that koalas throughout Victoria and SA are generally genetically similar to one another as their populations were mostly re-established from southern offshore islands at the beginning of the 20^th^ century, after they were earlier decimated by hunting [18–24]. This may have facilitated the dissemination of degraded KoRV and/or KoRV resistance in the south but could also directly explain the abrupt change in KoRV distributions between NSW and Victoria. Therefore, further investigation of KoRV’s genetic and geographic patterns is required to elucidate historic and contemporary KoRV-koala dynamics.

Retroviral subtypes are one important component of genetic diversity and can reflect host-virus evolutionary history and transmission dynamics. New variants/subtypes may be selected to allow superinfection in populations where virus incidence is high [25, 26]. KoRV can be delineated into several subtypes based on the amino acid sequence of the receptor binding domain of the envelope protein [27]. Only subtype A is known to occur endogenously [7, 28] and is also found as an exogenous virus throughout Australia [13, 14]. KoRV subtype B was first identified in a captive koala colony in Japan (referred to as KoRV-J) and was subsequently isolated from captive koalas in the San Diego zoo [3, 29]. It has since been detected in wild koalas throughout northern Australia [4, 5, 15, 30]. KoRV subtypes C, D, E and F were isolated from captive koalas internationally [29, 31], with KoRV-D subsequently found throughout wild northern koalas whereas KoRV-C has only been observed in northern Queensland and KoRV-F is more common in southern Queensland [15].

Chappell *et al.* [27] identified a further three subtypes (G-I) using deep sequencing in wild southern Queensland koalas and subtype K has recently been identified in captive Australian koalas [32]. Several of these exogenous subtypes have also been reported in southern koalas [16, 30, 33]. However, these results require confirmation as they were identified at around the detection limit for the methodology used (see discussion) and KoRV-B was not detected in southern Australia despite PCR screening of over 640 animals [6].

These recent studies suggest that KoRV subtype diversity is extensive, with some subtypes common throughout the koala’s range while others are more restricted. The processes by which these subtypes arise is currently unknown. It is likely that the hypervariable region of the receptor-binding domain is subject to positive selection for mutations and recombination events with host loci including other endogenous retroviral elements that can alter receptor use, and so overcome host cell resistance to super-infection, as shown to occur in endogenous feline leukaemia virus [34, 35]. In feline leukaemia virus, subtype A is the only subtype to be transmitted and undergoes recombination within each host to produce subtype B [36, 37]. However, KoRV subtypes appear to be exogenously transmitted predominately from mother to offspring [15, 32]. This suggests that rather than resulting from repeated recombination, each subtype may have arisen from a rare event and then spread through koala populations via exogenous transmission. Investigating the patterns of genetic diversity and differentiation across the koala’s geographic range should provide further insight into which of these processes is likely to predominate, with local differentiation among subtypes B to K in the absence of KoRV-A differentiation more consistent with exogenous transmission than repeated recombination.

In this study we analysed isolated DNA from washed koala faecal samples to describe KoRV genetic diversity across the koala’s natural range with the specific aims: (1) Confirm the geographic extent of KoRV endogenization and identify any populations where endogenization is not complete; (2) Characterise the geographic patterns and distributions of KoRV subtypes to gain insights into subtype evolution and transmission dynamics; and (3) Assess how KoRV genetic diversity both within and between populations varies over the koala’s geographic range to inform our understanding of KoRV evolution.

## Results

### KoRV incidence and copy number per cell

To investigate KoRV’s geographic patterns we successfully extracted DNA from 192 faecal wash samples collected from 20 populations throughout the entire koala geographic range of eastern Australia (Fig. 1A). All samples collected from NSW and QLD were KoRV *pol* positive as determined by qPCR (Fig 1B). By contrast, 25.8% (24 or 93) of samples collected from Victoria and SA were KoRV *pol* positive with individual population incidence rates ranging from zero to 55.5% (Fig 1B). Ulupna, situated on the Victorian side of the Murray River, which forms the NSW/Vic boarder, had the highest KoRV *pol* prevalence.

**Fig 1:**
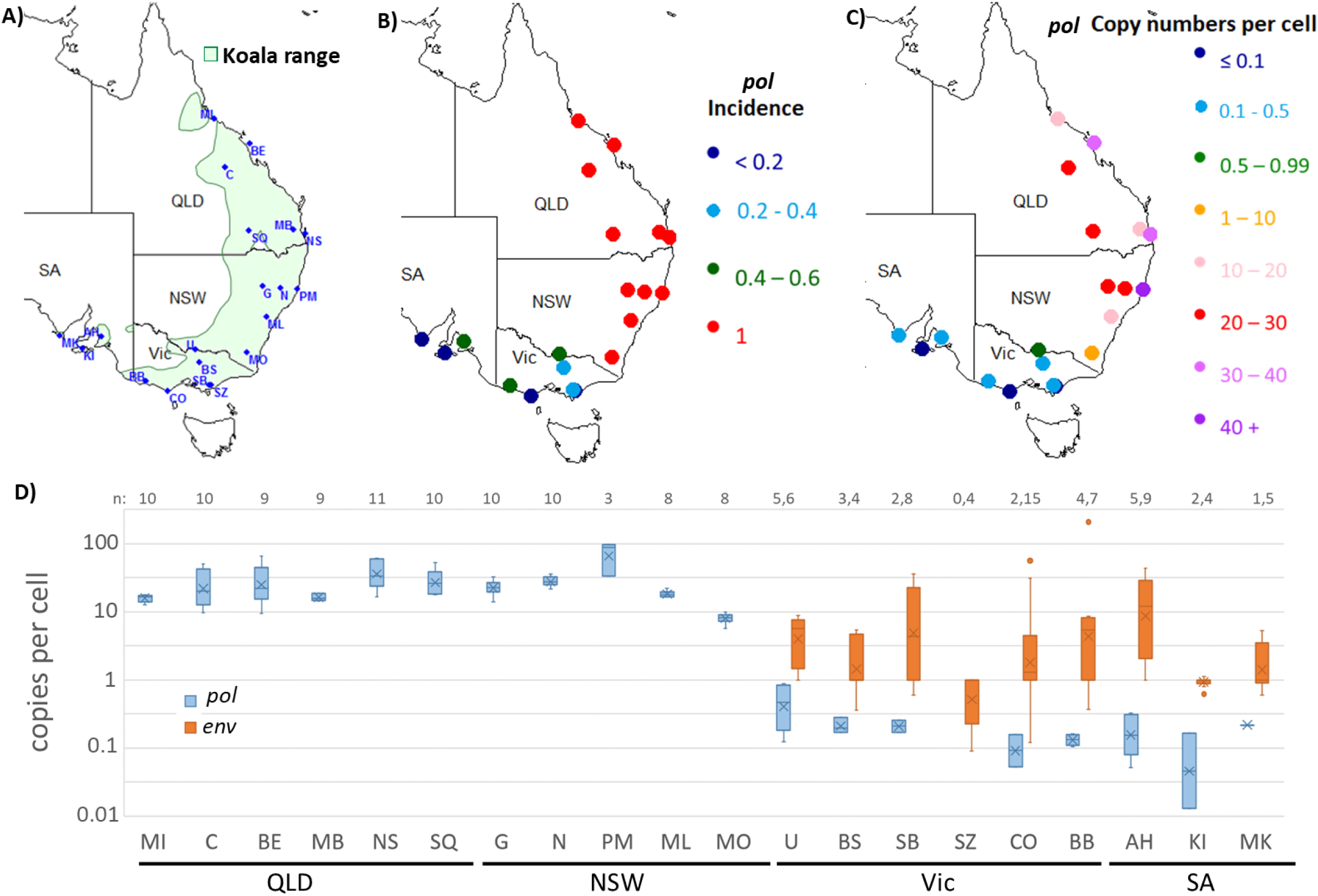
Geographic location (A), KoRV *pol* incidence (B) and average *pol* proviral copies per koala cell (C) for each population. (D) Boxplots of the number of KoRV *pol* and *env* copies per koala cell, values shown for *pol* positive koalas and southern koalas for which single *env* melt curve peaks were obtained, respectively. MI = Magnetic Island, C = Clermont, BE = St. Bee’s Island, MB = Mt. Byron, NS = North Stradbroke Island, SQ = South-West Queensland, G = Gunnedah, N = Nowendoc, PM = Port Macquarie, ML = Mountain Lagoon, MO = Monaro, U= Ulupna, BS = Boho South, SZ = Strzelecki 1, SB = Strzelecki 2, CO = Cape Otway, BB = Bessiebelle, AH = Adelaide Hills, KI = Kangaroo Island, MK – Mikkira . QLD = Queensland, NSW = New South Wales, Vic = Victoria, SA= South Australia

As identified previously [13], there was a distinct difference in KoRV *pol* proviral copy number per koala cell between the northern (QLD/NSW) and southern populations (Vic/SA). All northern koalas had estimated copy numbers greater than two, suggesting that KoRV was endogenous in those populations [38] (Fig. 1C-D). Copy number estimates varied considerably among northern koalas, ranging from 5.7 to 97.3 (Fig 1D). The lowest copy numbers were found in koalas from the Monaro region in southern NSW (mean = 8.1, n = 8) (Fig. 1D). Copy number estimates were considerably lower than the 133 previously estimated from examination of the genome of a northern koala [28] and the 139-199 previously estimated from blood/tissue samples [13]. However, our estimated copy numbers for Port Macquarie koalas were consistent with those previously reported for faecal samples collected from the same region [38]. Therefore, the variation in copy number between these studies may be due to differences in copy number between sample types due to reintegration of endogenous KoRV or exogenous infection in some tissues [39].

In the southern populations, estimated *pol* proviral copy number did not exceed one in any of the sampled koalas (Fig. 1D), which is inconsistent with the presence of proviral sequences in every cell as occurs when endogenous sequences are present. This strongly suggests that intact KoRV is not endogenized in those populations. However, it should be noted that these copy numbers are based on an estimate of 14 copies of β-actin in the koala diploid genome. If the actual number of β-actin copies was in fact 16 then the KoRV *pol* copy number in one of the Ulupna koalas would exceed one. Therefore, the presence of intact endogenous KoRV at low incidence should not be discounted for the Ulupna population.

In stark contrast to the findings for KoRV *pol,* the incidence of KoRV-A *env* among the 97 southern koalas was 100% as determined by qPCR. Of these koalas 35 produced melt curve peaks at multiple temperatures and 11 did not have a peak at the expected temperature for the original KoRV-A *env* fragment. This potentially indicates that these samples contained multiple sequence variants, preventing accurate copy number estimation. Of the remaining 62 samples, 46 (74.2 %) had greater than 1 *env* copy per cell (mean = 11.2, range = 1.1 – 56.3, excluding an outlier from Bessiebelle that had 209 copies per cell) (Fig 1D). This suggests that at least a fragment of the KoRV *env* gene is endogenous in the majority of southern koalas.

### Geographic distribution of KoRV subtypes

To determine KoRV genetic diversity and subtype incidence across the koala’s geographic range, Illumina deep sequencing was conducted on all KoRV *pol* positive samples (n=124) using an approximately 500 bp region of the KoRV *env* gene that includes the previously identified hypervariable region within the receptor-binding domain [27]. To confirm that faecal wash samples could be used to identify subtype presence, we sequenced paired blood and faecal wash samples from eight captive koalas. Subtypes were unable to be detected in the faeces when they occurred at less than 0.5% of *env* reads in plasma (n = 6). When subtypes were found above 0.5% abundance in the plasma, they were detected in the faecal washes on 65% (13/20) of occasions. Thus, in agreement with the findings of Quigley *et al.* [30], we determined that faecal wash samples provide a convenient and non-invasive tool for assessing KoRV subtype diversity, although, not all subtypes present, particularly those at low abundance, are reliably detected.

A total of 486 unique *env* sequences were characterised after *de novo* clustering of the quality controlled deep sequencing reads at 97 % similarity and removal of singletons. Of the unique sequences, 173 (35.6%) were found to contain deletions, frameshifts and/or nonsense mutations and were classified as non-functional. Among the remaining 313 intact sequences, the previously identified subtypes A (n = 87), B (n = 19), C (n = 15), D (n = 118), F (n=1), I (n = 8), and K (n =3) were detected from a combination of PCoA analysis (Tables 1 and 2; Fig S1), nucleotide similarities (NTS) of the hypervariable region and the *in silico* translated amino acid sequences of the hypervariable region. Subtypes E (12), G, and H (2) were not found. Among the subtype D sequences, the PCoA analysis revealed 39 sequences that were not tightly clustered with the other D sequences and many of those had low similarities to the D reference sequence (minimum 70.1% NTS). However, the mutations in these sequences were consistent with stepwise mutation from the remaining tightly clustered D sequences and showed amino acid homology to the reference D sequence. Therefore, they were designated ‘divergent D’ sequences. The translated amino acid sequences also revealed several intermediates between subtypes, with the first section of the hypervariable region showing homology to one subtype and the remainder resembling a different subtype. These sequences included 17 A/B intermediates, 8 A/D intermediates and 4 D/F intermediates (Table 2). Two distinct sequence clusters that did not show similarity to any of the previously identified subtypes (Fig S2 and S3) and were designated subtypes L and M. The hypervariable region of subtype L showed some similarity to subtype F (71.6 to 82.0% NTS), while subtype M showed low similarity to all previously identified subtypes (maximum NTS 63.3 to 72.9%).

**Table 1:** Incidence of KoRV subtypes among koala populations

| **Subtype** | **Total number of**<br>**koalas** | **Number of**<br>**populations** |
| --- | --- | --- |
| A/B intermediate | 28 | 7 |
| B | 12 | 3 |
| C | 9 | 3 |
| D | 60 | 10 |
| divergent D | 19 | 7 |
| A/D intermediate | 10 | 4 |
| F | 2 | 1 |
| D/F intermediate | 6 | 3 |
| I | 4 | 1 |
| K | 1 | 1 |
| L | 6 | 1 |
| M | 4 | 1 |

**Table 2:**
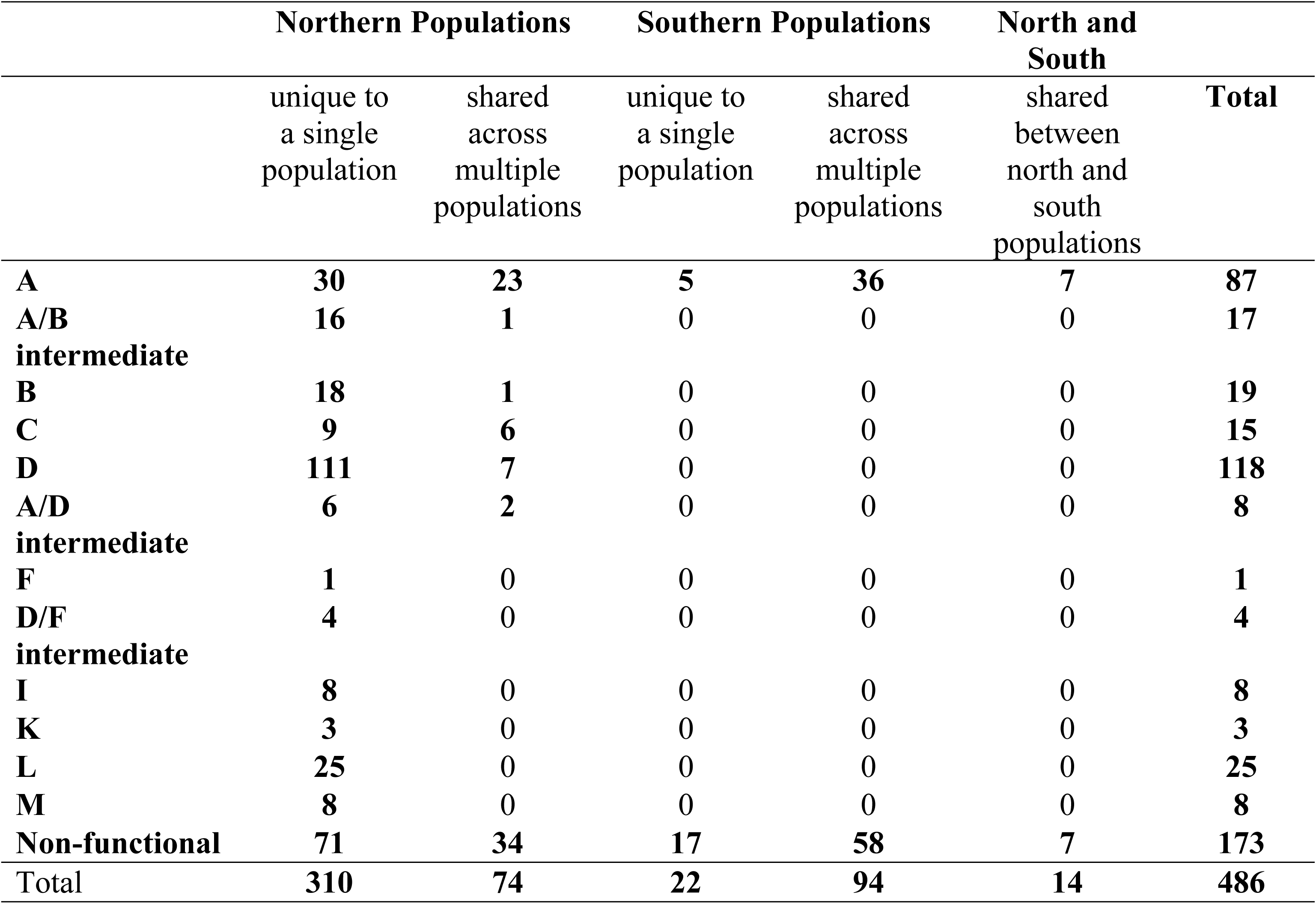
Geographic distribution of KoRV sequence clusters

In northern koala populations, KoRV-A was the dominant subtype and found in all koalas, with abundance ranging between 83.5 to 100% of KoRV reads. In southern populations, KoRV-A was the only subtype detected (n = 24). The predominant KoRV-A sequence present in all animals clustered with 97% similarity to the original KoRV-A sequence isolated by Hanger *et al.* [40] and later confirmed to be endogenous in northern koalas [7]. The majority (77.4%) of koalas also possessed additional sequences belonging to KoRV-A at low frequency (maximum 2.1%). Non-functional sequences were also detected in most (87.1%) koalas but at very low combined frequency (average = 0.4%, maximum = 5.2%).

To further investigate KoRV-A sequence similarity among koalas, the sequencing reads were re-clustered at 100% similarity. This revealed four sequences that each accounted for the majority of reads in one or more koalas. One of these sequences was the original KoRV-A sequence and was found in all northern koalas and 8 southern koalas. This sequence predominated in NSW and Southern Queensland koalas but was generally at low abundance in North Queensland (Fig 2). Another sequence that differed from the original sequence by one nucleotide was abundant among Queensland koalas but rare in NSW. A third sequence that also differed from the original KoRV-A by one nucleotide was most abundant in north NSW. Among southern *pol* positive koalas, all except one animal from the Adelaide Hills had the same dominant sequence that differed by three nucleotides from the original KoRV-A (Fig 2).

**Fig 2:**
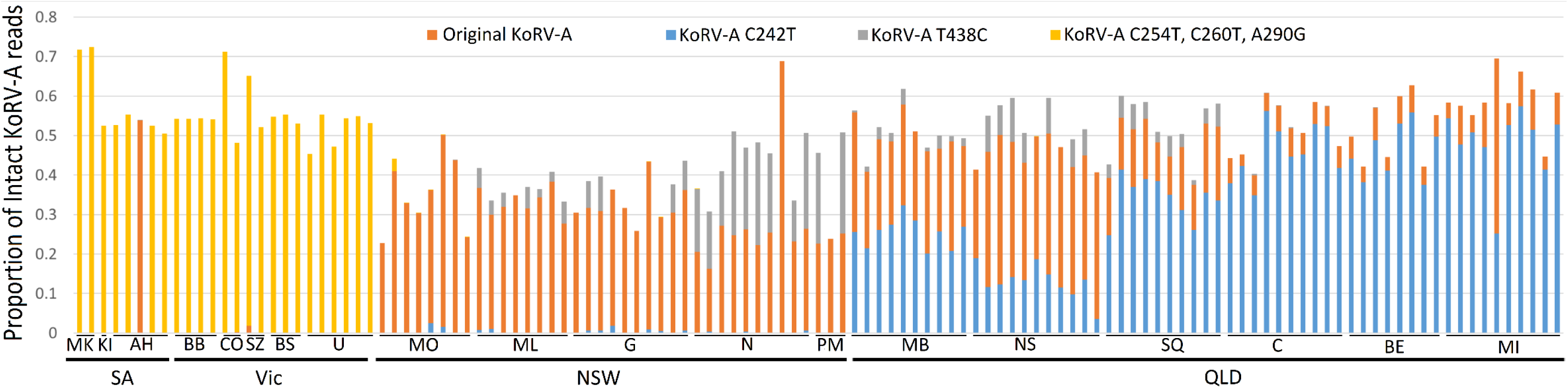
Proportion of intact KoRV-A reads from each KoRV *pol* positive koala belonging to the four dominant sequences. Mutation codes are relative to the original KoRV-A sequence [40] with numbering relative to the beginning of the *env* gene. Population and State codes as described in Figure 1.

Among the northern koalas, KoRV-D was the most prevalent non-KoRV-A subtype and was detected in 60.6% (60/99) of koalas that were widely distributed across 10 of the 11 populations (Table 1; Fig 3B). Divergent D sequences and A/D intermediate sequences were also widely distributed across multiple populations, with divergent D sequences also found in a population where D was not detected (Fig 3B). A/B intermediate sequences were more prevalent than KoRV-B sequences and were found in 7 populations, including 3 in NSW where KoRV-B has not been detected (Fig 3A). Different A/B intermediate sequences were detected in each population, however, the breakpoints between the A and B motifs were in similar positions among the majority of sequences (Fig S4). By contrast, KoRV-B was only recovered from 3 QLD populations. Subtype C was restricted to northern QLD, while subtypes I, L and M were locally prevalent (40%, 75% and 50%, respectively) but each restricted to a single sampled population (Fig 3C). Subtype F itself was only found in two koalas from Mt. Byron (QLD) (Fig. 3C). However, D/F intermediates reached an incidence of 36.4% (4/11) on North Stradbroke Island and were found in a single koala in two other populations. Subtype K was only detected in a single koala from northern QLD (Fig 3C).

**Fig 3:**
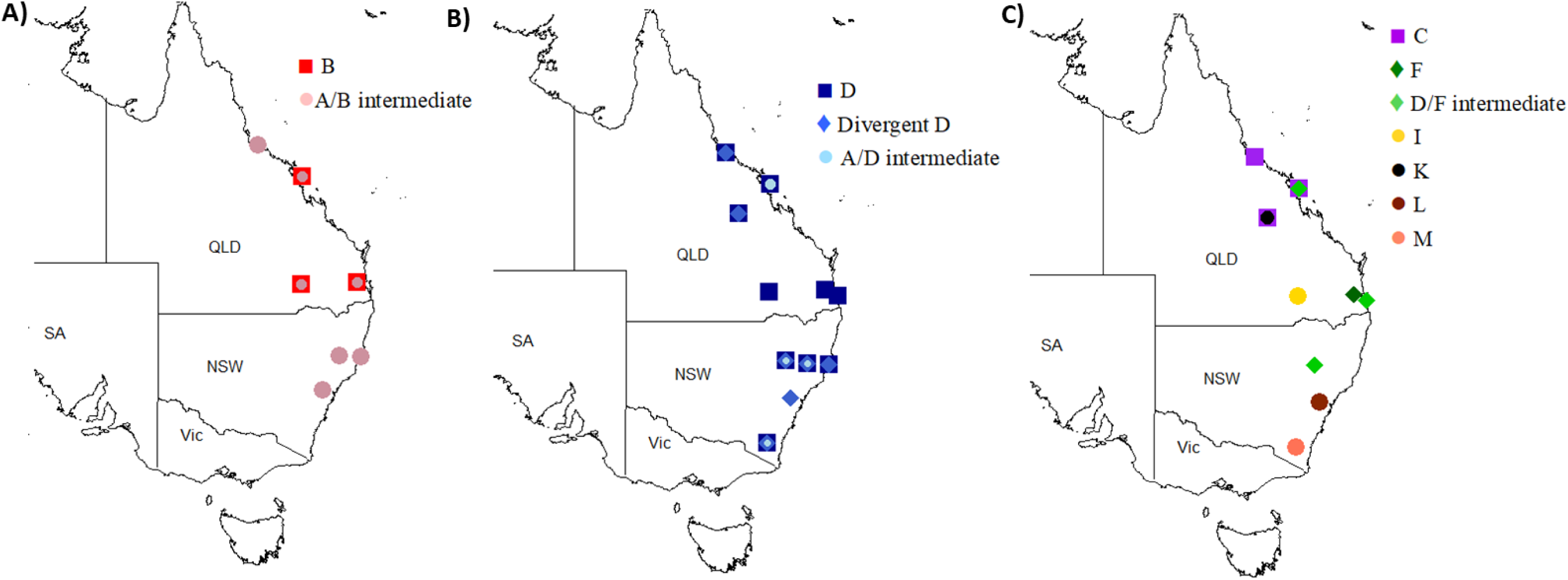
Geographic distribution of KoRV-B and A/B intermediates (A); KoRV-D and A/D intermediates (B); and the other detected subtypes (C)

### env sequence sharing and richness

Of the 486 unique *env* sequence clusters 312 (64.2%) were detected in multiple koalas, 162 (33.3%) were shared between populations and only 14 were shared between northern and southern koalas (Table 2). Among these sequences, 384 were detected in northern koalas and 116 were detected in southern koalas. Similarly, among the 313 intact sequences there were far more identified in northern koalas (279 sequence clusters) than among southern koalas (41 sequence clusters) with only a very small number (7 sequence clusters) shared between the north and the south (Table 2). This was not an effect of differing sample sizes between regions as shown by rarefaction (Fig. 4D).

**Fig 4:**
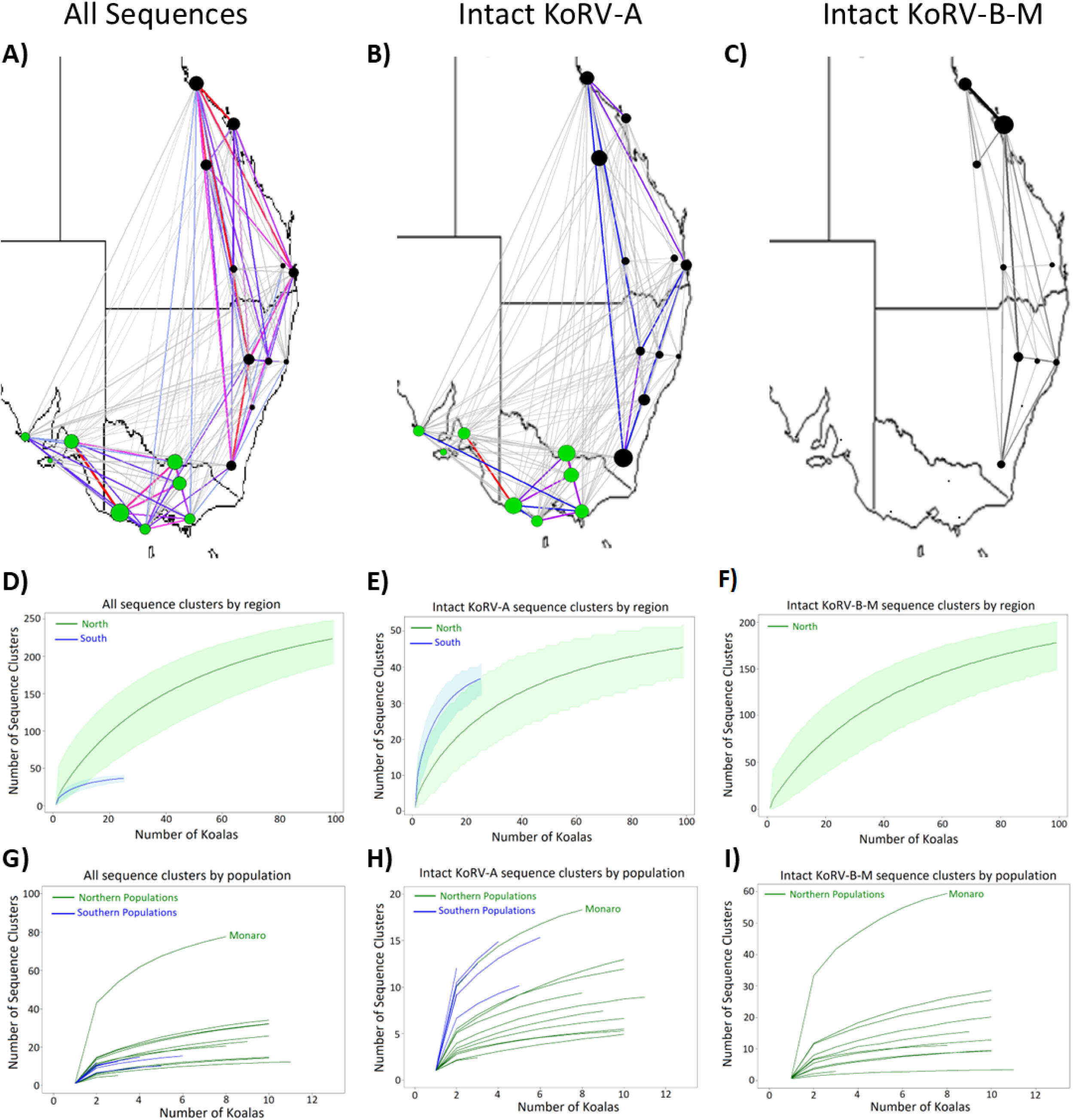
*Env* sequence sharing and richness. Network diagrams of all (intact and non-functional) KoRV (A), intact KoRV-A (excluding the original KoRV-A cluster) (B) and intact KoRV-B-M sequence clusters shared among populations. Nodes represent sites, with the size of the node indicating the number of sequence clusters shared with other populations. Black nodes = northern populations, light green nodes = southern populations. Lines between nodes represent shared sequence clusters, with the thickness of the line indicative of the number of sequence clusters shared. Grey lines = 5 or less sequence clusters shared; gradient of blue to red lines = 6-20 clusters shared. Rarefaction curves of the total number of intact KoRV sequences (D and G), intact KoRV-A sequence clusters (E and H) and intact KoRV-B-M sequence clusters (F and I) detected with increasing sample size across (D, E and F) and within (G, H and I) northern and southern populations. 95% confidence intervals are shown for D, E and F.

At the population level, there was considerably less sequence cluster sharing between northern and southern populations than among populations within those regions (average number of shared sequence clusters between two populations: north vs south = 1.57, north vs north = 6.78, south vs south = 7.68; Fig 4A). However, within the north or south there was no evidence that populations that were geographically closer together shared more sequence clusters (Mantel test with significance determined by permutation, north: p = 0.18; south: p = 0.13).

The number of intact sequences found within populations for the same number of sampled koalas was generally similar in the north and south (Fig. 4G). This suggests that the high total number of unique sequences in the north is due to differences between populations in the sequences found, rather than higher within population richness (Table 2). An exception to this general finding is the Monaro population (NSW), where a higher number of sequence clusters were identified with each additional koala sampled than for the other populations (Fig 4G).

The patterns of intact KoRV-A sequence sharing among populations reflected those seen for all sequences combined, with less sharing between the north and the south than within those regions (Table 2, Fig. 4B). However, in contrast to the patterns observed for all intact sequences combined, a similar number of unique KoRV-A sequence clusters were identified among southern (n =38) and northern (n = 40) populations, despite fewer koalas being sampled in the south (Fig. 4E). This indicates that each southern koala carries more unique KoRV-A sequences than do northern koalas and this is reflected in a higher number of unique sequences being identified per southern population, after accounting for differences in sample sizes (Fig. 4H). Again, the Monaro population was found to be an exception and had similar KoRV-A richness to the southern populations (Fig 4H).

Compared to KoRV-A there were far fewer KoRV-B-M sequences shared among populations (KoRV-A = 59.8%, KoRV-B-M = 7.5%; Fig. 4B-C, Table 2). KoRV-B to M richness could not be compared among regions due to the absence of the exogenous subtypes in the south. However, among northern populations, the Monaro population was again an outlier and had far more unique KoRV-B to M sequences than the other populations (Fig. 4I).

### env genetic diversity throughout the koala range

In order to assess genetic diversity in the *env* sequence without the added complexity of subtype presence, the pairwise number of nucleotide differences between sequences was calculated, excluding the hypervariable region (that is likely to be derived through recombination and thereby does not adhere to stepwise diversification). In general, the mean number of nucleotide differences (MND) among all pairs of intact and non-functional sequence clusters detected within a population was significantly higher for the northern populations than for the southern populations, although not by a large margin (mean ± SD: north = 17.35 ± 3.03, south = 13.13 ± 2.83; p =0.007; Fig 5A).

**Fig 5:**
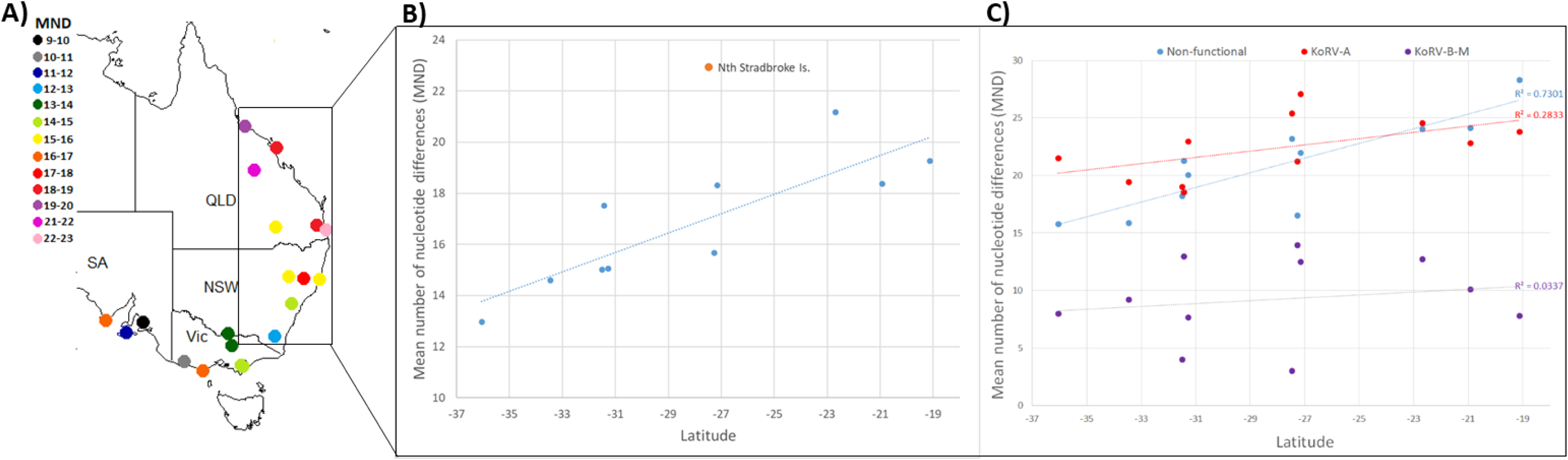
The mean number of pairwise nucleotide differences among all sequences detected within each population by geographic location (A) and for northern populations in relation to latitude (B). Dotted line = fitted regression, blue points = populations included in regression analysis, orange point = Nth Stradbroke Island. The mean number of pairwise nucleotide differences among KoRV-A, KoRV-B-M and non-functional sequences (C) are also shown in relation to latitude for the northern populations.

There was no distinct geographic pattern in sequence diversity for southern populations (Fig. 5A). Whereas sequence diversity (MND) within the northern populations significantly decreased with distance from the equator (R^2^ = 0.48, p = 0.018). The naturally occurring koala population (not introduced) on North Stradbroke Island had higher sequence diversity than expected from its latitude and the strength of the correlation between MND and latitude was considerably improved by its exclusion (R^2^ = 0.72, p = 0.001; Fig 5B). When considering the different types of sequence clusters separately, non-functional sequence diversity was strongly correlated with latitude (R^2^ = 0.73, p < 0.001; Fig 5C). There was a trend for intact KoRV-A *env* diversity to increase with proximity to the equator (R^2^ = 0.28, p = 0.09), but there was no evidence of a correlation between latitude and sequence diversity among subtypes B to M (R^2^ = 0.03, p = 0.58; Fig 5C).

The ‘genetic centre’ of all populations appears to be the original KoRV-A sequence as it had the lowest MND to other sequences in each population. The Monaro was the only exception, with a divergent D sequence and the KoRV-A original sequence having similar MND (8.72 and 8.74, reactively). Once the hypervariable region was excluded, 18 sequences belonging to subtypes B (n = 1), C (n = 2), D (n = 8), divergent D (n = 3), D/F intermediate (n = 1), I (n = 1) and L (n = 1) were identical to the original KoRV-A sequence. Additionally, the MND among KoRV-B to M sequences was far lower than among KoRV-A sequences across all populations, suggesting they have more recently diversified (Fig 5C).

### env sequence genetic differentiation

Analysis of molecular variance (AMOVA) based on the diversity of sequences (the pairwise nucleotide differences among the sequence clusters) in each population revealed small but significant KoRV *env* genetic differentiation between northern and southern koalas (variance attributed to differences between regions = 2%; p =0.004; Table 3). There was no differentiation between southern populations (variance attributed to differences between southern populations = 0 %; p = 0.755), while there was significant differentiation accounting for 8% of the total genetic variance across northern populations (p =0.001; Table 3). How different the northern populations were from one another varied between pairs of northern populations (Table S1), however, populations that were geographically more distant were not more genetically different (Mantel test between geographic distance and PhiPT: p = 0.208; Fig S5). Instead, the diversity of KoRV sequences within each northern population was distinct (Table S1), with the Monaro considerably more distinct than the other populations (average pairwise PhiPT: Monaro = 0.14; other populations = 0.03 to 0.08).

**Table 3:** Summary of molecular variance among populations within regions and between regions

|  | Between | Between | Between |
| --- | --- | --- | --- |
|  | Northern populations | Southern populations | North and South |
| Variation between populations by sequence presence only |  |  |  |
| All sequences | 8% (p=0.001) | 0% (p=0.755) | 2% (p=0.004) |
| KoRV-A | 1% (p =0.18) | 0% (p = 0.993) | 4% (p<0.001) |
| KoRV-B-M | 19% (p < 0.001) | NA | NA |
| Non-Functional | 3% (p=0.001) | 0% (p = 0.859) | 3% (p<0.001) |
| Variation between populations by sequence and incidence |  |  |  |
| All sequences | 10% (p=0.001) | 1% (p = 0.012) | 2% (p = 0.001) |
| Variation between populations by sequence, incidence and abundance |  |  |  |
| All sequences | 26% (p=0.001) | 0% (p = 0.995) | 26% (p = 0.001) |

When considering the different types of sequence clusters separately, KoRV-A differentiation was non-significant among northern and southern populations, while there was a low but significant level of differentiation among regions (Table 3). There was also no differentiation among southern populations for the non-functional sequence clusters, while there was low but significant differentiation among northern populations (Table 3). By contrast, there was considerable differentiation in KoRV-B to M subtype sequence clusters between northern populations (despite the exclusion of the hypervariable region from the analysis; variance attributed = 19%; p < 0.001; Table 3). Thus, the total *env* genetic differentiation observed between northern populations can primarily be attributed to KoRV-B to M.

### The effect of population incidence and within koala abundance on genetic differentiation

When the incidence of (number of koalas carrying) the sequence clusters within each population was included in the AMOVA analysis (instead of only their presence), overall differentiation among northern populations increased slightly from 8% to 10%, while very low but significant differentiation was found among southern populations (Table 3). Further, when the sequences were also weighted by their average within koala abundance in the AMOVA, differentiation among northern populations increased markedly to 26%, while there remained no differentiation among southern populations (Table 3). This marked increase in differentiation among northern populations reflects the trend for the more abundant sequence clusters within koalas to be found in fewer populations (regression z value = -1.753, p = 0.080; Fig S6b).

Among sequence clusters detected in northern koalas, those that had a high incidence within a population were also found at higher abundance within koalas from that population (regression z value = 3.277, p = 0.001 Fig S6a). Therefore, between-koala differences in KoRV *env* sequence composition that accounted for 7% of genetic variation based on incidence (p= 0.001) became non-significant when within koala abundance was taken into account (genetic variance = 1%, p =0.554; Fig. 6). These locally abundant sequence clusters predominantly belonged to KoRV-B to M, with only 2 of the 13 most prevalent sequence clusters within northern populations found to belong to KoRV-A (after exclusion of the ubiquitous original KoRV-A cluster).

**Fig 6:**
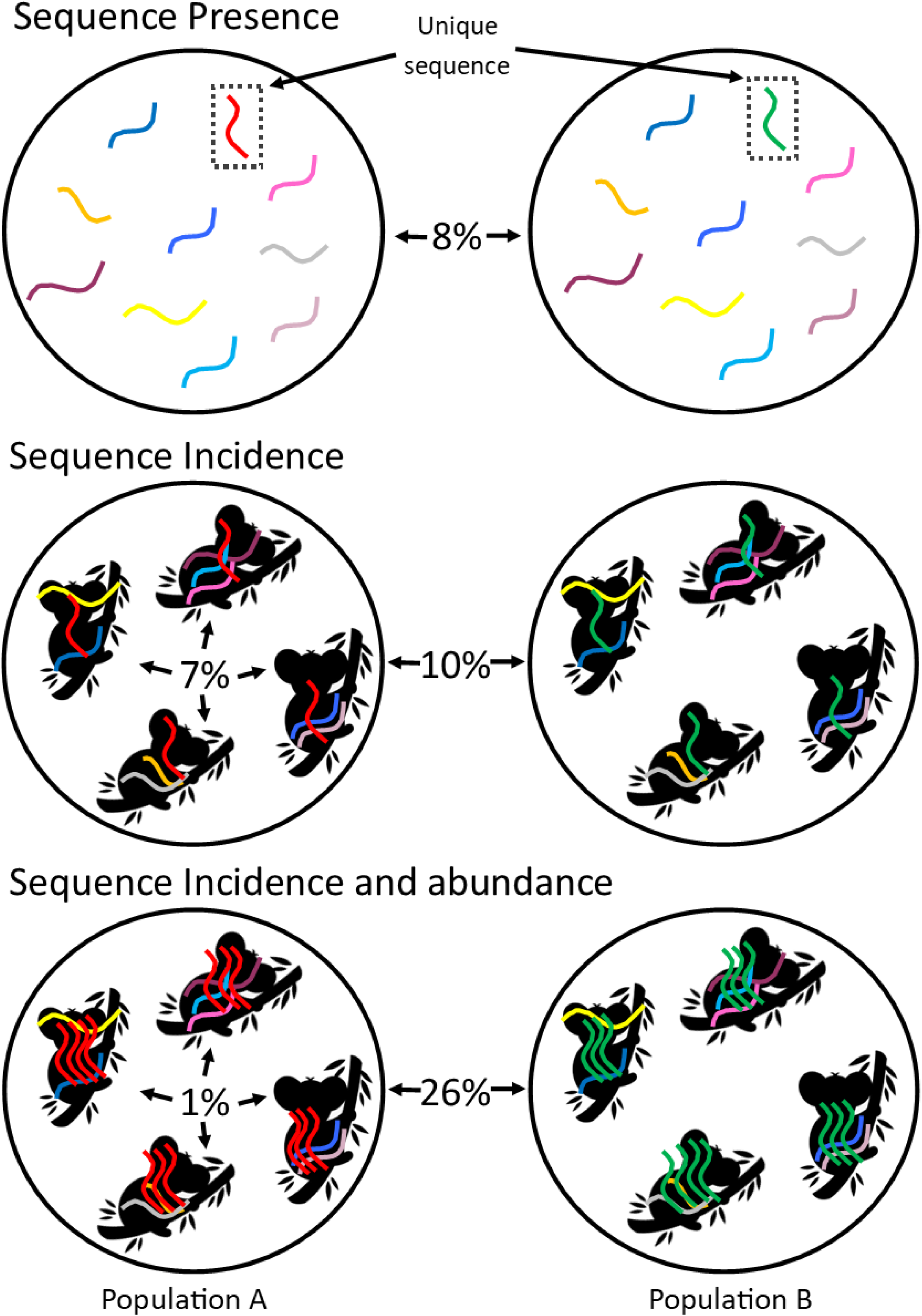
Diagram of how genetic differentiation between **northern populations** is influenced by sequence incidence and within koala abundance as reflected by the percentages of molecular variance generated from the AMOVAs.

## Discussion

Our assessment of the biogeographic distribution of KoRV sequence diversity provides new insights into KoRV evolution, the processes underlying the emergence of novel subtypes and has implications for koala conservation. Here we discuss each of these topics in turn and outline how our findings challenge several previous assumptions to contribute to an improved understanding of KoRV.

### KoRV evolution

In agreement with previous studies [6, 13, 14, 30], our *pol* gene incidence and copy number findings show that all koalas in the northern Australian states of QLD and NSW appear to carry intact, endogenous KoRV. However, in contrast to assumptions made in previous studies based solely on *pol* gene data [13, 38] or KoRV-A specific *env* PCR [30], our results suggest that the majority, and potentially all, southern koalas carry partial KoRV-A sequences within their genomes. While partial sequences consisting of the terminal ends of the KoRV genome, including partial *env*, have previously been identified [16, 17], our work shows that such sequences are widespread and additionally that all southern koalas carry an identical *env* sequence which is distinct from the originally described KoRV-A sequence [40] at three nucleotides.

The presence of partial endogenous KoRV sequences across southern koalas but not endogenous full-length KoRV-A has implications for the resounding question of how and where KoRV evolved. Given the relatively young age of KoRV-A (less than 50,000 years (2)), the successful elimination of all full-length copies of KoRV from the genomes of southern koalas is unlikely, even when considering the severe population bottleneck that has occurred in southern populations (13, 23). Alternatively, higher KoRV sequence diversity in northern Queensland is consistent with a longer history of KoRV infection (Fig7) and suggests that intact KoRV-A first entered the koala genome in northern Australia. It is therefore possible that full length KoRV-A may not have yet spread to southern koalas or it may be present in only a very low number of animals. However, if this is the case then it raises the question of how partial replication incompetent endogenous KoRV sequences became ubiquitous in southern koalas in the absence of replication competent KoRV.

**Fig 7:**
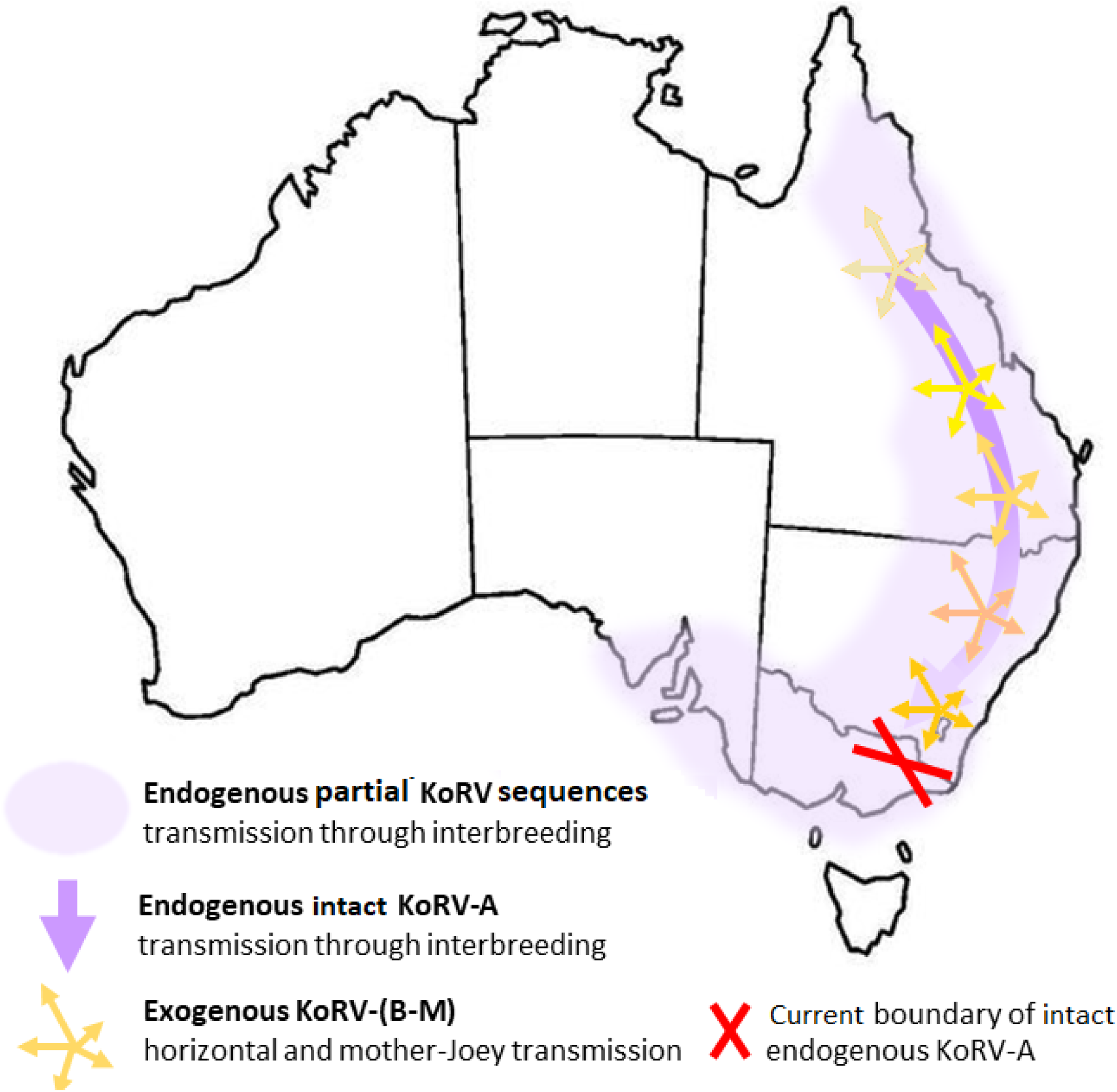
Schematic of proposed KoRV evolution hypothesis

Further research is required to determine how and when these partial KoRV sequences invaded southern koala populations, though it is likely that it involved other endogenous and exogenous viral elements. Löber *et al.* [17] identified partial KoRV sequences in the koala genome flanking a partial sequence of an older degraded retroelement (PhERV) and named these sequences recKoRV based on their hypothesis that such sequences were the result of recombination with intact KoRV-A. However, analysis of KoRV-A sequence insertions within the reference koala genome reveals a clear difference between those insertions comprising truncated KoRV LTRs (12-35% divergence) and those of full length KoRV-A (<2% divergence) (Fig. S7), suggesting that the truncated sequences may be considerably older and may pre-date the emergence of full-length KoRV-A. Therefore, the genealogical relationship among the recKoRV sequences, partial KoRV sequences and full-length KoRV-A is not clear. Simmons et al (39) identified a similar virus to KoRV in an Australian native rodent (*Melomys burtoni*) and other similar viruses have recently been detected in flying foxes (40, 41). While none of these viruses are the proximate source of KoRV, they demonstrate that multiple closely related retroviruses exist and that there is a wealth of viral biodiversity that may have contributed to contemporary patterns of both full length and partial KoRV sequences.

In addition to host-virus co-evolutionary processes, it appears that koala population history and demography has impacted the geographic distribution of KoRV. Our assessment of proviral copy numbers, subtype profiles and KoRV genetic diversity all showed abrupt changes at the Victoria/NSW border. Koalas in the Ulupna population on the Victorian boarder had low genetic diversity and were genetically similar to other koalas in Victoria, while the genetic diversity of the Monaro population in southern NSW was consistent with that of other NSW koalas (unpublished microsatellite data). This is consistent with the re-introduction of koalas throughout Victoria and SA from southern off-shore islands [13, 23]. This historic population bottleneck can explain the absence of KoRV genetic differentiation and the high rate of sequence cluster sharing (despite small sample sizes) among southern populations. Among northern koalas, KoRV *pol* copy number, genetic diversity and subtype profiles in the Monaro population were found to be divergent from the other populations. Recent phylogenetic analysis of koala populations throughout Australia has revealed a shallow lineage division around the Sydney harbour basin in central NSW, which may indicate a past biogeographic barrier in that region [22]. Such a barrier could have reduced the transmission of both exogenous and endogenous KoRV between the Monaro population and the rest of northern Australia, leading to the observed KoRV differentiation.

### The emergence of novel subtypes

We did not detect any non-KoRV-A subtypes in any of the koalas we sampled from southern Australia (Victoria and SA). This is consistent with previous PCR based analyses that have not detected KoRV-B in Victoria [6] but is at odds with the results of two deep sequencing studies of koalas in the Mt. Lofty Ranges, South Australia [16, 33] and one of koalas in Victoria [30]. The first of the Mt. Lofty Ranges studies [16] inferred the presence of the non-KoRV-A subtypes from the pseudo-alignment of a very small number of reads to the hypervariable regions of the different subtypes. Given the small read counts, these assignments are likely to be erroneous and our re-analysis of this data (ENA: PRJEB21505) using competitive read mapping revealed no compelling evidence for the presence of non-KoRV-A subtypes in their southern koalas (results not shown). In the second study focusing on koalas within the Mt. Lofty Ranges study [33], there was remarkable similarity between the subtype sequences detected in their Queensland and South Australian study sites, with 93% of sequences shared and an 80% correlation between the number of koalas carrying each sequence in the two populations (calculated from the supplementary information). Further, all non-KoRV-A sequences identified in the Victorian koalas were also found in northern koalas from that study [suplimentry information; 30]. Given that the sequence sharing between two adjacent Queensland captive populations that exchange koalas was only 46% [Joyce personal communication; 32], high sequence sharing between northern and southern populations would be unexpected and it is more likely that the low abundance detection of the non-KoRV-A subtypes in the Mt. Lofty Ranges and Victorian studies were due to cross contamination during sample preparation or sequencing.

Our findings suggest that new exogenous KoRV subtypes arise from endogenous KoRV-A under selection to escape host suppression and allow superinfection where functional KoRV-A is ubiquitous. In all populations (with the exception of the Monaro) the original endogenous KoRV-A sequence cluster was the genetic centre of sequence diversity, with the representative sequences from clusters of six other subtypes found to be identical to that sequence after the hypervariable region was excluded. Additionally, the other subtypes were only detected in populations where intact KoRV-A was endogenous. These findings on their own could lend support to the hypothesis that KoRV-A repeatedly gives rise to the other subtypes within hosts, as occurs in Feline Leukaemia Virus [36, 37]. However, we found that several of the subtypes had localised geographic distributions, with considerable KoRV genetic differentiation between northern populations due to the non-KoRV-A subtypes. This finding suggests that different subtypes arise sporadically across the geographic range and become locally prevalent through selection and exogenous transmission (Fig 7), which is known to occur at least between mothers and their offspring [15, 32].

### Implications for koala conservation

The koala is now listed as vulnerable by the Australian Federal Government and faces serve decline in north-eastern Australia due to habitat loss and disease [41, 42]. The characterisation of KoRV subtype diversity across the koala’s range will likely contribute to the management of disease in the future, particularly for northern koala populations. Little is currently known about the disease association of the majority of the KoRV subtypes. However, there is some evidence that KoRV-B may be more pathogenic than KoRV-A as it has been associated with Chlamydiosis and neoplasia in southern Queensland koalas [3–5]. Therefore, it is of concern that KoRV-B has recently been detected in NSW and Queensland by deep sequencing [15, 30] and that we have also identified KoRV-B in northern Queensland and A/B intermediates in NSW. As unique A/B intermediates were detected in each population they may have been produced by local hybridisation instead of widespread transmission of intermediate sequences. Alternatively, since directionality of recombination cannot be assumed and the A/B intermediates were more widely detected than KoRV-B, these intermediate sequences may in fact be ancestral and KoRV-B is the product of two recombination events. Further investigation of the clinical significance of all the subtypes is needed to more robustly ascertain how their distributions are likely to impact koala health in different areas of Australia. However, it is clear that as the exogenous subtypes are currently geographically restricted, the translocation of koalas among northern populations would lead to the introduction of novel subtypes to a region, with potentially negative impacts on disease.

Whether replication competent, non-germline integrated KoRV is present in southern koalas is an important question with implications for how southern populations are managed. In the south, our estimated number of *pol* copies per cell were markedly lower than one and were detected in around 26% of animals. Such findings have previously been interpreted as the presence of functional exogenous KoRV in some southern koalas [13, 38], however it should be noted that the full length KoRV genome has never been identified in southern koala populations. Nevertheless, a previous study in over 640 koalas did find an association between KoRV-A *pol* positivity and poor body condition in Victorian koalas [6], with a significance level of p= 0.008, suggestive of the presence of replication competent KoRV-A. Studies to determine the presence of the entire functional KoRV genome in the south should be high priority to determine whether functional exogenous KoRV is a threat to koalas in the south.

The stark difference in KoRV incidence and diversity between northern and southern koala populations highlights priorities for management and potential containment of KoRV. In a recent study investigating the transmission of exogenous KoRV-sequences amongst captive northern koala populations [32], we found low sequence sharing between non-related, co-housed animals and no evidence of sexual transmission. Transmission of exogenous KoRV sequences was overwhelmingly from mother to offspring. Unless an alternate mode of transmission is at play within southern koalas, neither the exogenous subtypes nor endogenous KoRV-A are anticipated to spread into the south quickly and translocations between northern and southern populations are not advised. Instead, koala populations at the boundary of the northern and southern koala populations (around the NSW/Vic border) are of high interest. The Monaro population in southern NSW displays low proviral copy numbers [30, this study] and yet high sequence richness. This could be attributed to more recent endogenization and a greater expression of KoRV due to an absence of strong host suppression. Additionally, the presence of koalas with *pol* proviral copy numbers approaching 1 in the Ulupna population could be the result of endogenous functional KoRV-A recently reaching the border through migrant koalas from southern NSW.

## Methods

### Study sites and sample collection

Koala faecal samples were collected between April and September 2016, from 20 locations (Fig. 1) that span the koala’s current wild geographic range. The habitats at the sites varied between cleared farmland with semi-isolated paddock trees, open woodland and tall eucalypt forest. In general, faecal samples were collected from 8-11 individuals per site with the exception of Port Macquarie (NSW) where three animals were sampled and Cape Otway (Victoria) where 18 koalas were sampled. Fresh faecal pellets were collected from each koala by placing a plastic mat (2 m by 3 m) directly beneath them. Faecal pellets were collected from the mat and immediately frozen at -18 °C. All samples were collected less than 24 hours post-production and 98 % were collected less than 10 hours after production. Latitude and longitude were recorded at the location of each sampled koala using a Garmin eTrex 20x handheld GPS. These coordinates were used to calculate the geographic centre of each study site.

To determine how KoRV subtype profiles recovered from faecal DNA compared to those obtained from plasma RNA, paired blood and faecal samples were collected from eight captive koalas housed at Currumbin Wildlife Sanctuary, Dreamworld on the Gold Coast and Paradise Country Wildlife Park. Up to 3 mls of blood was drawn by a qualified veterinarian from the cephalic vein of each koala during conscious restraint. The blood was then transferred to an EDTA coated tube to prevent clotting. Faecal samples were collected concurrently directly from the koalas’ cloacas into sterile 15 ml tubes. The samples were stored on ice for approximately 3 to 5 hours during transport to the laboratory. The blood was then centrifuged at 16 000 g for 10 minutes and the plasma collected by pipetting. The plasma and faeces were then stored at -80 °C.

The fieldwork was carried out under Western Sydney University ethic approval (A11253) and with appropriate permits from the New South Wales (SL101722), Queensland (WITK17277716), Victorian (10007714) and South Australian State governments (U26533-1). Collection of samples from captive koalas was carried out under University of Queensland ethic approval (AE36153).

### Molecular techniques

#### Nucleic acid Extraction

Total DNA was extracted from surface washes of the faecal samples following the general approach of Wedrowicz et al. [43]. A single faecal pellet from each koala was placed in 3 mls of 1% phosphate buffered saline and gently rotated for 15 minutes on a rotatory mixer. The pellet was then removed and discarded. The phosphate buffered saline wash was centrifuged for 15 minutes at 4 000 g and the supernatant removed. DNA was extracted from the wash pellet by Digsol/proteinase K digestion following the method of Bruford *et al.* [44] and eluted in 50 ul of 1x Tris-EDTA. PCR inhibitors were then removed from the extracted DNA using the OneStep PCR inhibitor removal kit (Zymo) following the manufacturer’s instructions.

Prior to RNA extraction the plasma samples were passed through a 0.45 µm membrane filter to remove any remaining koala cells. Total RNA was extracted from the plasma samples using the High Pure Viral Nucleic Acid Kit (roche) according to the manufacturer’s instructions with modification as outlined in the supplementary methods. First strand cDNA was then created from the RNA extracts using Superscript III (Invitrogen) and 32 µM random pentadecemer primers [45] following the manufacturer’s instructions. Following synthesis, the cDNA was treated with 1 ul of RNase H (New England Biolabs) and incubated at 37 C of 20 minutes.

#### Quantitative PCR

The presence of the KoRV *pol* gene in a faecal sample was inferred from the successful amplification of a 110 bp fragment by quantitative PCR (qPCR) run in triplicate using previously designed primers [46]. The presence of the KoRV *env* gene was inferred from the amplification of a 97 bp (in the case of KoRV-A) fragment using degenerate primers designed from the alignment of KoRV-A and KoRV-D *env* sequences [27] (KoRV-RBD(A,D).F : CTCACTGCAWCGGCCTCYCAACAGGC; KoRV-RBD(A,D).R: GGGATAGCTACATCCCAGGGTTYC). A 123 bp fragment of the koala β-actin gene was also amplified in triplicate as a positive control [46]. Each fragment was amplified in a 10 µl reaction containing 1X SYBR™ Green PCR Master Mix, each primer at 500 nM and 2 µl of template. Tenfold dilutions of the DNA were first used as template, with the amplification repeated with 100-fold dilutions for those samples that showed evidence of inhibition or that failed to amplify for β-actin.

Amplification was performed using a QuantStudio 6 Flex Real-Time PCR system using the standard comparative CT protocol with an annealing temperature of 60°C. Quality control and standard curves were then completed as per the supplementary methods.

#### env hypervariable domain sequencing

An approximately 500 bp region of the KoRV *env* gene containing the previously identified hypervariable domain was amplified and sequenced from all KoRV positive samples (identified by qPCR). Of these samples, the *env* hypervariable region could be amplified in 64 using the protocol of Chappell et al. [27], with the number of PCR cycles varied between 28 and 40 depending on KoRV DNA quantity. Adequate amplification could only be obtained for the remaining 57 faecal wash samples and the plasma cDNA using the nested polymerase chain reaction (PCR) protocol described in the supplementary methods. For the faecal DNA, either 10 (n =91) or 100 (n = 30) fold dilutions were used as PCR template to overcome the presence of any remaining PCR inhibitors in the extractions.

Sequencing libraries were prepared from the *env* PCR products using the Nextera XT sample preparation kit (Illumina) following the manufacturer’s protocol. Each library was uniquely tagged using the Nextera XT indexing primers (Illumina). Paired-end sequencing was performed at the Australian Centre of Ecogenomics, on the Illumina Nextseq using the version 2 reagent kit for 300 cycles.

### Bioinformatic processing and statistical analysis

#### KoRV incidence and copy number per cell

KoRV incidence was inferred from the proportion of β-actin positive samples that were also positive for KoRV *pol* or *env* as determined by qPCR. The number of KoRV gene copies per koala cell was estimated for each sample from the number of KoRV *pol* or *env* and β-actin molecules per qPCR reaction. The number of KoRV *pol, env* and β-actin molecules present in each reaction were calculated from the average CT values and standard curves using the appropriate standard equations. To estimate the number of β-actin copies per koala cell, the number of fragments produced from the koala genome (NCBI assembly: GCA_900166895.1) with the qPCR primers was determined in CLC genomics workbench 20. This analysis produced 7 fragments or 14 copies per diploid cell. Therefore, the KoRV gene copies per cell were estimated as: 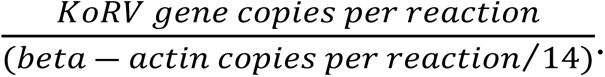

#### Geographic distribution of KoRV subtypes

The geographic distribution of KoRV subtypes was determined from the *env* Illumina deep sequencing data. The raw reads were merged, quality filtered and rarefied to 10 000 reads per sample in CLC genomics workbench 20 as described in the supplementary methods. The rarefied reads were then *de novo* clustered across all samples at 97% similarity in QIIME 2 [47]. Clusters containing only a single read across the dataset were discarded. To determine if the representative KoRV sequences were putatively functional, they were translated *in silico* with those containing missense and frameshift mutations or large deletions designated as non-functional as described in the supplementary methods. Subtype designations for the intact representative KoRV sequences were assigned by comparison to a set of reference subtype sequences in a three-step process as described in the supplementary methods. The geographic distribution of the subtypes was then inferred from the representative sequence by sample table generated from QIIME 2 clustering.

#### env sequence richness, genetic diversity and differentiation

To assess how the genetic diversity of KoRV varied within and among populations over the koala’s geographic range both the intact and non-functional representative KoRV *env* sequences were analysed to capture the complete KoRV “genetic pool” within which recombination is presumed to be prolific. Sequence incidence and within koala abundances were calculated from the representative sequence by sample table generated from the QIIME 2 clustering. Network diagrams of sequence sharing were constructed in NetDraw. Rarefaction curves and 95% confidence intervals of the number of unique sequences detected with increasing sampling effort were constructed using the package ‘rich’ [48] in R [49].

When calculating the sequence similarities, the hypervariable portion of the *env* nucleotide sequences (corresponding to aa81-143 of KoRV A numbering) was excluded as it may be of non-KoRV origin in some subtypes [34]. The pairwise number of nucleotide differences between sequences were calculated in GenAlEx 6.5 [50, 51]. Analysis of Molecular Variance (AMOVA) and pairwise PhiPT values were also calculated using GenAlEx 6.5 [50, 51] with significance determined by 9999 permutations of the dataset. The AMOVA was carried out using three different underlying versions of the data. In the first analysis, each sequence detected in a population was included once in the distance matrix, with sequences found in multiple populations included once for each population. This analysis tested sequence differentiation among populations. In the second analysis, the incidence of each sequence within a population was accounted for by including each sequence found within a koala once (e.g., if a sequence was detected in 5 koalas in a population it was included in the distance matrix 5 times for that population). This analysis also measured variation between koalas within populations in the sequences they carried. In the third analysis, the values in the distance matrix from the second analysis were weighted by the within koala relative abundance of the sequences (multiplied by 1000). Thus, this analysis accounted for within koala relative abundance and incidence. However, the weighting of the distance matrix reduces the validity of the permutation test of significance and thus significance values associated with this analysis should be viewed with caution.

The geographic distance between populations was calculated from the decimal latitude and longitude values for the centre of each site. Mantle tests were then performed in GenAlEx to test for associations between geographic and genetic distances. Generalised and linear regression models were fitted with the stats package in R using appropriate link functions. Regression models that included random effects were fitted using the lme4 package [52] in R with significance determined using the lmerTest package [53].

## Acknowledgements

The following people assisted with study site selection and fieldwork: Mark Boorman, Bill Ellis, Sean FitzGibbon, Ben Barth, Dalene Adam, Huiying Wu, Cheyne Flanagan, Mathew Crowther, Caroline Marschner, Laura Schmertmann, Daniel Lunney, Mark Krockenberger, Valentina Mella, Kellie Leigh, Faye Wedrowicz, James Fitzgerald, Kath Handasyde, Jack Pascoe, Emily Hynes, Robyn Molsher and Jason VanWeenen. We thank Peter Smouse for his advice on the statistical analysis of KoRV genetic diversity and differentiation.

## Declarations

The fieldwork was funded through a grant from the Australian Research Council (LP140100751) in partnership with Evolva Biotech A/S, the NSW Office of Environment and Heritage and The Conservation Ecology Trust. The molecular analysis was funded by the Australian Research Council (DP180103362).

